# Single-Cell Analysis of *NF1*-Expressing and *NF1*-Deficient Schwann and Fibroblast Cells Reveals Divergent Neurofibroma Programs

**DOI:** 10.1101/2025.09.26.677858

**Authors:** Grace M Swanson, Marcia Arenas-Hernandez, Katherine Gurdziel

## Abstract

Neurofibromatosis type 1 (NF1) is an inherited tumor predisposition syndrome characterized by the development of benign peripheral nerve sheath tumors, most commonly cutaneous neurofibromas (cNFs) and plexiform neurofibromas (pNFs). Although both tumor types arise from Schwann cells and share *NF1* loss as a genetic driver, they differ markedly in growth behavior, microenvironmental context, and clinical outcomes, with pNFs carrying risk of malignant transformation. To define transcriptional programs that underlie these differences, we performed an integrative single-cell RNA sequencing analysis of Schwann cells and fibroblasts from cNF and pNF tumors, alongside *NF1*-expressing reference populations derived from human skin and nerve. This approach enabled us to disentangle *NF1*-dependent transcriptional changes from tissue-of-origin effects. We identified distinct Schwann cell states that separated tumor from non-tumor populations and further distinguished cNFs from pNFs, highlighting both shared disease-associated features and subtype-specific adaptations. These findings establish a framework for understanding how *NF1* loss interacts with developmental origin and tissue context to shape divergent tumor phenotypes and may inform strategies for therapeutic targeting in NF1-associated neurofibromas.

## 1. Introduction

Neurofibromatosis type 1 (NF1) is an inherited tumor predisposition syndrome caused by biallelic inactivation of the tumor suppressor gene *NF1*, which encodes the Ras-GTPase activating protein neurofibromin a negative regulator of Ras signaling (1, 2). Loss of NF1 function promotes hyperactivation of multiple pathways including Ras/MAPK, PI3K/Akt/mTOR, ROCK/LIMK/cofilin, and cAMP/PKA, with downstream effects on cellular proliferation, migration, cytoskeletal remodeling, and differentiation (3–7). Clinical manifestations of NF1 are highly variable and affect multiple organ systems, but nearly all patients develop benign peripheral nerve sheath tumors known as neurofibromas (8). Neurofibromas are broadly categorized into cutaneous neurofibromas (cNFs), which arise in the dermis, and plexiform neurofibromas (pNFs), which extend along large nerve plexuses and can infiltrate adjacent tissues (1). While cNFs remain benign and are not associated with metastatic progression, pNFs carry a 10–15% lifetime risk of malignant transformation into malignant peripheral nerve sheath tumors (MPNSTs), the most aggressive and life-threatening complication of NF1 (9, 10). Despite originating from Schwann cells and sharing a common genetic etiology, cNFs and pNFs exhibit strikingly different growth behavior, microenvironmental context, and clinical outcomes, suggesting the involvement of distinct downstream molecular programs.

In addition to their shared genetic origin, cNFs and pNFs exhibit overlapping histological and transcriptional profiles, complicating efforts to define tumor subtype-specific disease mechanisms (8, 11). Bulk transcriptomic analyses have identified gene expression signatures associated with neurofibroma subtype, but these are often confounded by differences in cellular composition, anatomical location, and inter-individual variability (12, 13). Moreover, both cNFs and pNFs contain a complex mixture of Schwann cells, fibroblasts, immune cells, and vascular cells, making it difficult to disentangle the cell-intrinsic consequences of *NF1* loss from microenvironmental influences (14–17). Single-cell RNA sequencing (scRNA-seq) provides an opportunity to overcome these limitations by resolving transcriptomic heterogeneity at cellular resolution, enabling a more precise definition of disease-relevant cell states and signaling programs (18–20). Furthermore, the ability to compare tumors arising in different anatomical contexts from the same individual offers a powerful strategy to identify subtype-specific features while controlling for inter-individual genetic background.

In order to gain insight into the molecular mechanisms that distinguish cutaneous and plexiform neurofibromas, we performed an integrative analysis of single-cell transcriptomic datasets with a focus on Schwann cells and fibroblasts since they are the primary cellular constituents of these tumors. In the current study, our goal was to identify transcriptional programs associated with *NF1* gene loss that may contribute to tumor subtype identity, development and behavior. To achieve this, we utilized two complementary single-cell RNA sequencing (scRNA-seq) datasets derived from either neurofibroma (*NF1*-deficent) or non-tumor (*NF1*-expressing) matched tissues within individual donors. The first dataset, published by Wang et al. (11), includes scRNA-seq profiles of cNF and pNF resected from the same patient with neurofibroma type 1 (NF1), enabling a direct comparison of tumor phenotypes in a shared genetic context. The second dataset, published by Chu et al. (21), consisted of Schwann cells and fibroblasts derived from non-tumoral human skin and nerve tissue paired samples from 3 individuals; therefore, providing a reference for tissue-specific transcriptional baselines. These two datasets offer a unique opportunity to disentangle the effects of tissue context from tumor-specific transcriptional alterations, and to identify transcriptional drives that may underlie the divergent trajectories of cNF and pNF development.

To define the transcriptional consequences of *NF1* gene loss in distinct tumor contexts, we performed an integrative scRNA-seq analysis of Schwann cells and fibroblasts derived from neurofibromas (cNF or pNF respectively), and non-tumoral skin and nerve tissue. Cell-type-specific clusters were identified across datasets, and differential gene expression and pathway enrichment analyses were conducted to uncover shared and divergent molecular programs. By resolving tumor- and non-tumoral-tissue-specific programs at cellular resolution, the work herein provides new insight into the mechanisms that distinguish cNF and pNF and establishes a framework for identifying potential therapeutic targets in *NF1*-deficent tumors.

## 2. Materials and Methods

### 2.1 Data sources and sample processing

Single-cell RNA sequencing datasets were obtained from Gene Expression Omnibus (GEO): GSE218493 (Wang et al.) (11) and GSE125422 (Chu et al.) (21). In total, eleven samples were analyzed, two samples from dataset GSE218493 containing one cNF and one pNF, both derived from the same individual with neurofibroma type 1 (NF1). From dataset GSE125422, we analyzed nine non-tumoral tissue–derived samples from individuals expressing *NF1*, consisting of three biological replicates each of skin-derived Schwann cells, nerve-derived Schwann cells, and skin-derived fibroblasts. These datasets were selected because they provide paired tumor and non-tumoral samples from matched anatomical locations and shared donor backgrounds, enabling comparative analysis of cell type–specific transcriptional programs in the context of *NF1* gene loss.

Briefly, tumor tissues from Wang et al. (GSE218493) were obtained from a single individual with NF1 and processed through mechanical dissociation and enzymatic digestion in PBS containing 0.5% collagenase type I and 2% fetal bovine serum at 37 °C for 3 h. Cell suspensions were filtered, erythrocytes lysed using ACK buffer, and viable cells isolated by exclusion of 7-aminoactinomycin D–positive cells via fluorescence-activated cell sorting. scRNA-seq libraries were generated with the 10x Genomics Chromium platform (11).

Schwann cell and fibroblast datasets from Chu et al. (GSE125422) were generated from non-tumoral human skin and sciatic nerve tissues obtained from adult donors through the Southern Alberta Organ Donation Program or elective biopsies. Tissues were dissociated with dispase and collagenase, then Schwann cells were isolated using antibodies against the p75 neurotrophin receptor, and fibroblasts were isolated using antibodies targeting Thy-1. Cells were expanded in culture media supplemented with forskolin and neuregulin-1 prior to scRNA-seq library preparation on the 10x Genomics Chromium platform (21).

### 2.2 Bioinformatic analysis

#### 2.2.1 UMAP integration and cluster identification

The compiled scRNA-seq datasets were evaluated using Seurat in R (4.4.1) (22). Datasets were imported as individual Seurat objects, with Chu et al. neurofibroma tumoral samples mapped onto the Wang et al. reference non-tumoral samples to enable joint clustering and visualization. For Wang dataset, quality control retained cells with <10% mitochondrial gene content and 200– 20,000 detected features. Data were normalized and scaled before principal component analysis (PCA), and the top 14 principal components were selected for downstream analysis.

For Chu dataset, quality control followed the published workflow retaining cells with 1,500–6,000 detected genes and <4.5–7.5% mitochondrial RNA. Dataset integration was performed using Seurat’s canonical correlation analysis (CCA)-based workflow. To enable direct comparison of tumor- and non-tumoral-derived cell populations, we defined the Wang et al. tumor samples as the reference dataset since it contained multiple cell types and the Chu et al. non-tumoral tissue samples consisting of Schwann and fibroblast cells as the query dataset. Anchors were identified using the FindTransferAnchors function (CCA, default parameters) to map shared features between datasets. The MapQuery function was then used to project the non-tumoral tissue cells into the low-dimensional space defined by the integrated neurofibroma tumoral cells, ensuring that Schwann and fibroblast populations from *NF1*-expressing skin and nerve aligned with their tumor-derived counterparts.

Cluster identification was conducted using shared nearest neighbor (SNN) graph–based clustering, and dimensionality reduction was visualized using uniform manifold approximation and projection (UMAP). Conserved markers across samples and differentially expressed genes (DEGs) between clusters were identified using Seurat’s FindConservedMarkers and FindMarkers functions. DEGs were considered significant with an adjusted *p*-value < 0.05 and absolute log2 fold change > 0.25. Cell types were annotated primarily by manual curation using canonical marker genes from the literature; predictions from ScType were used as a secondary check (23).

#### 2.2.2 Subsetting and reclustering of Schwann cells and fibroblasts

Following global integration, Schwann cell and fibroblast populations were identified by expression of canonical markers (*SOX10*, *S100B*, *NGFR* for Schwann cells; *VIM*, *PDGFRA*, *DCN* for fibroblasts) and subset from the integrated Seurat object. For fibroblast-barcoded cells, likely Schwann cell contaminants were removed if they met either of the following criteria: location within the Schwann manifold (UMAP x∈[−15,−9], y∈[0,5]) or expression of ≥ 3 Schwann markers (*SOX10, ERBB3, S100B, PRX, MPZ, PMP2, PMP22, PLP1, NRXN1, GALR1, XKR4, PRIMA1, COL28A1, SOX2*). Each subset was then reprocessed independently using the same Seurat workflow: normalization (NormalizeData), scaling (ScaleData), dimensionality reduction by principal component analysis (PCA), and reclustering with a shared nearest neighbor (SNN) graph-based approach (FindNeighbors, FindClusters). Dimensionality reduction and visualization of subclusters were performed using UMAP (RunUMAP).

Differential expression (DE) analyses were performed within each lineage using the FindMarkers function (Wilcoxon rank-sum test, default parameters). Genes were considered significantly differentially expressed at an adjusted *p*-value < 0.05 and absolute log2 fold change > 0.25.

#### 2.2.2 Functional enrichment analysis

Functional enrichment was performed using the clusterProfiler R package (24). Differentially expressed genes (DEGs) from each condition were analyzed using the enrichGO, enrichKEGG, and enrichPathway functions to identify significantly enriched Gene Ontology (GO), KEGG, and Reactome terms, respectively. Enrichment results were compiled across conditions and converted into data frames for downstream analysis in R.

For GO enrichment, terms were further collapsed into higher-level functional categories using a combination of manual curation (mapping specific GO identifiers to defined bins) and keyword-based classification of term descriptions. Broad functional groups included mitochondrial metabolism, RNA processing, protein homeostasis, immune response, extracellular matrix and adhesion, and developmental signaling.

For each condition and category, enrichment strength was summarized by the maximum and median −log10 adjusted *p*-value, and the number of terms contributing to each bin was recorded. Results were visualized in ggplot2 as heatmaps (enrichment strength as color, term counts overlaid as labels) and dot plots (dot size proportional to term count, color scaled by enrichment significance).

#### 2.2.3 Receptor-ligand binding analysis

Receptor-ligand binding analysis was performed for signals coming from either Schwann cell or fibroblast clusters using CellChat in R (25). Interactions were inferred from Schwann cell and fibroblast clusters directed toward all other identified cell types. Four major interaction modes were evaluated: secreted signaling, extracellular matrix (ECM)–receptor interactions, cell–cell contact, and non-protein interactions. CellChat analyses were conducted independently for cutaneous neurofibroma (cNF) and plexiform neurofibroma (pNF). Results were interpreted in the context of differentially expressed genes to highlight tumor-specific signaling events.

#### 2.2.3 Gene Ontology (GO) and upstream regulator analysis

To identify biological processes enriched in differentially expressed genes, we performed Gene Ontology (GO) enrichment analysis using the enrichGO function from the clusterProfiler R package (v4.6.2) (26). Differentially expressed genes (DEGs) were identified using the Seurat FindMarkers function with default Wilcoxon rank-sum testing. Genes were filtered for significance based on adjusted *p*-value < 0.05 and log2 fold-change thresholds specific to each comparison.

## 3. Results

### 3.1 Integrated Single-Cell Analysis Defines Tumor- and Subtype-Specific Cell States in Neurofibromas

To investigate how *NF1* gene loss alters Schwann cell and fibroblast transcriptional programs in neurofibromas, we performed an integrative single-cell RNA-seq analysis of cNF and pNF tumors together with non-tumoral skin- and nerve-derived Schwann cells and fibroblasts (11, 21). This design enabled cell type–specific comparisons between tumoral and non-tumoral contexts while controlling for tissue origin. We identified transcriptionally distinct Schwann cell states that not only segregated tumoral from non-tumoral populations but also revealed reproducible subtype-specific differences between cNF and pNF.

From the cNF and pNF samples, we identified 10 cell populations from 20 distinct transcriptional clusters (Figure 1A, Supplemental Figure 1A) (11). From the 20 clusters, cluster 13 was removed by filtering low quality cells. Inspection of canonical marker expression (Figure 1B, Supplemental Table S1) confirmed that the transcriptional clusters from the integrated neurofibroma dataset corresponded to the 7 major cell populations previously defined by Wang et al. (Schwann cell, fibroblast, smooth muscle, endothelial, perineurial/keratinocytes, macrophage, T-cell plus perivascular, lymphatic endothelium, and melanocytes). In addition to the characteristic markers used for cell-type assignment, the dot plot highlights distinctive lineage-enriched transcripts: Schwann cell (*S100B, PMP2, NRXN1*), perivascular (*S100B, PMP2, NRXN1, LUM, PDGFRA, OGN*), fibroblast (*LUM, PDGFRA, OGN*), smooth muscle (*RGS5, GUCY1A2, MRVI1, ADRA2A, CARMN*), endothelial (*MECOM, PTPRB, SELE*), lymphatic endothelium (*LYVE1, MMRN1, CCL21*), perineurial (*KRT15, DSG3, CXADR*), macrophages (*MS4A7, CD86, IL1B, CCL22*), melanocytes (*TRPM1, DCT, TYRP1, SLC6A15*), and T cells (*CD3D, CD3G, IL7R, IKZF3*). These features complement canonical lineage markers and emphasize cell type–specific transcriptional programs that distinguish stromal, vascular, immune, and neural lineages within the neurofibroma microenvironment.

**Figure 1.**
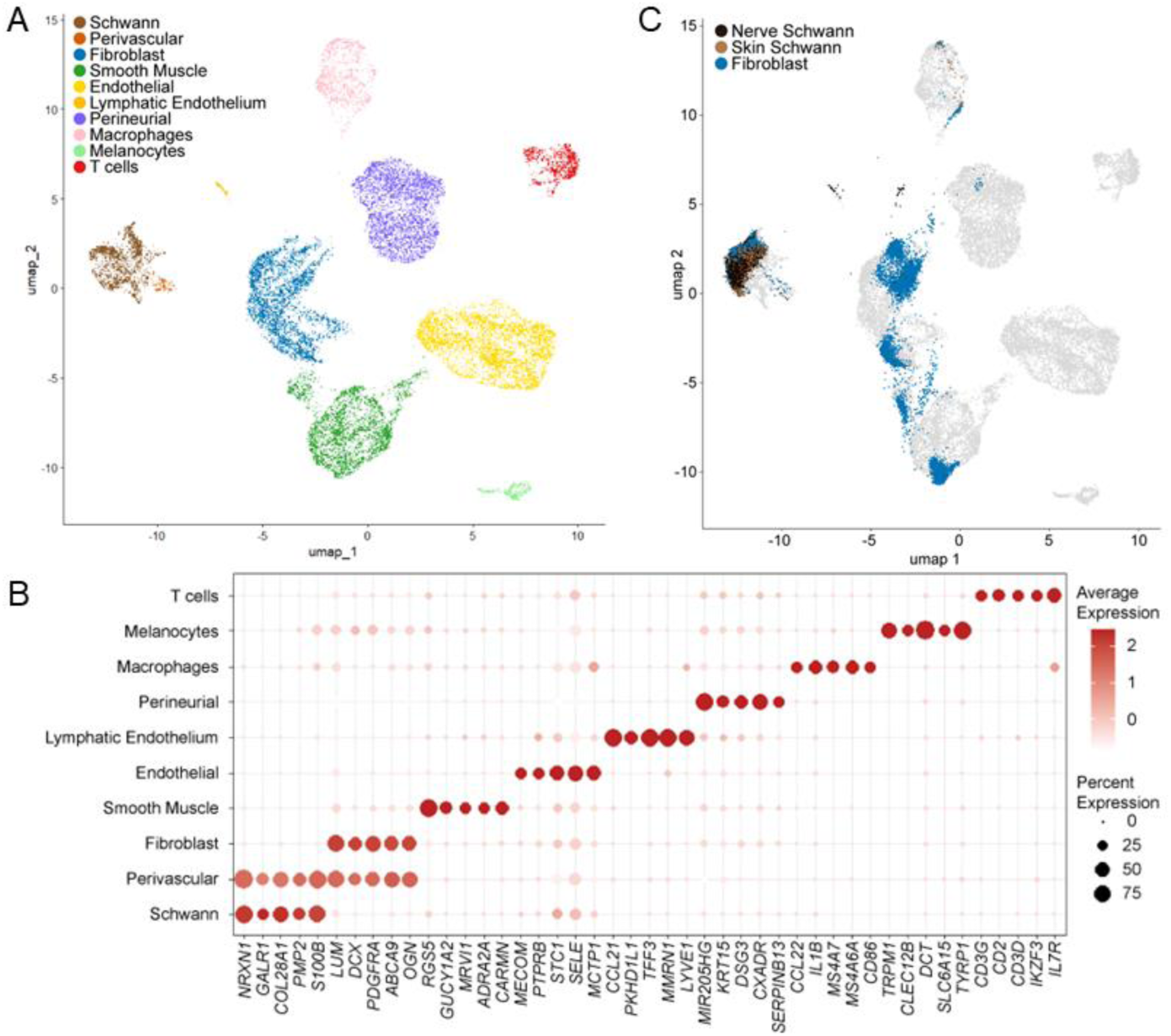
Identification and characterization of major cell populations across integrated single-cell datasets. **(A)** UMAP embedding of integrated cutaneous neurofibroma, plexiform neurofibroma, and *NF1*-expressing reference datasets, showing major cell types identified by canonical markers, including Schwann cells, fibroblasts, perivascular cells, smooth muscle, endothelial cells, lymphatic endothelium, perineurial cells, macrophages, melanocytes, and T cells. **(B)** Dot plot of selected marker genes used for cell type annotation. Dot size represents the proportion of cells expressing each gene within a given cluster, while color intensity reflects average expression level. **(C)** Projection of NF1-*expressing* reference populations (skin-derived Schwann cells, nerve-derived Schwann cells, and skin fibroblasts) into the integrated UMAP space, demonstrating alignment with their respective tumor-associated states.

### 3.2 Cross-Dataset Projection Reveals *NF1* Status and Tissue-of-Origin Differences in Schwann and Fibroblast Lineages

To compare *NF1*-deficient tumor-derived populations with *NF1*-expressing counterparts, we next projected Schwann cells and fibroblasts from the Chu non-tumoral dataset onto the Wang neurofibroma (tumoral) reference atlas (Figure 1C). *NF1*-expressing skin-derived Schwann cells aligned primarily with the Schwann cell clusters identified in the *NF1*-deficient neurofibromas, while *NF1*-expressing nerve-derived Schwann cells mapped to a distinct subset; therefore, highlighting tissue-of-origin differences. Similarly, *NF1*-expressing dermal fibroblasts projected onto the *NF1*-deficent fibroblast populations of the neurofibroma dataset, confirming lineage identity while revealing tumor-associated transcriptional divergence. This cross-dataset projection demonstrates that the 10 neurofibroma cell populations encompass both canonical lineages and N*F1*-gene loss–associated adaptations, providing a framework for direct comparison between *NF1*-expressing and *NF1*-deficient states.

After defining the major cell populations in the neurofibroma dataset, we next compared their distribution within cNF and pNF tumors. Both tumor types contained the ten populations identified above (Figure 1AB, Figure 2A), but their relative contributions differed markedly (Figure 2B). cNF were enriched for perineurial and T cell populations, compared to pNFs which displayed a lower proportion of the latter yet higher proportion of Schwann cells, fibroblasts and perivascular cells. Skin- and nerve-derived Schwann cell overlap revealed origin-specific variation (Figure 2C), with skin-derived cells localizing near cNF-associated Schwann states and nerve-derived cells mapping more closely to pNF-associated states. These patterns indicate that both neurofibroma tumor subtype and tissue of origin shape the transcriptional programs of *NF1*-deficient Schwann cells.

**Figure 2.**
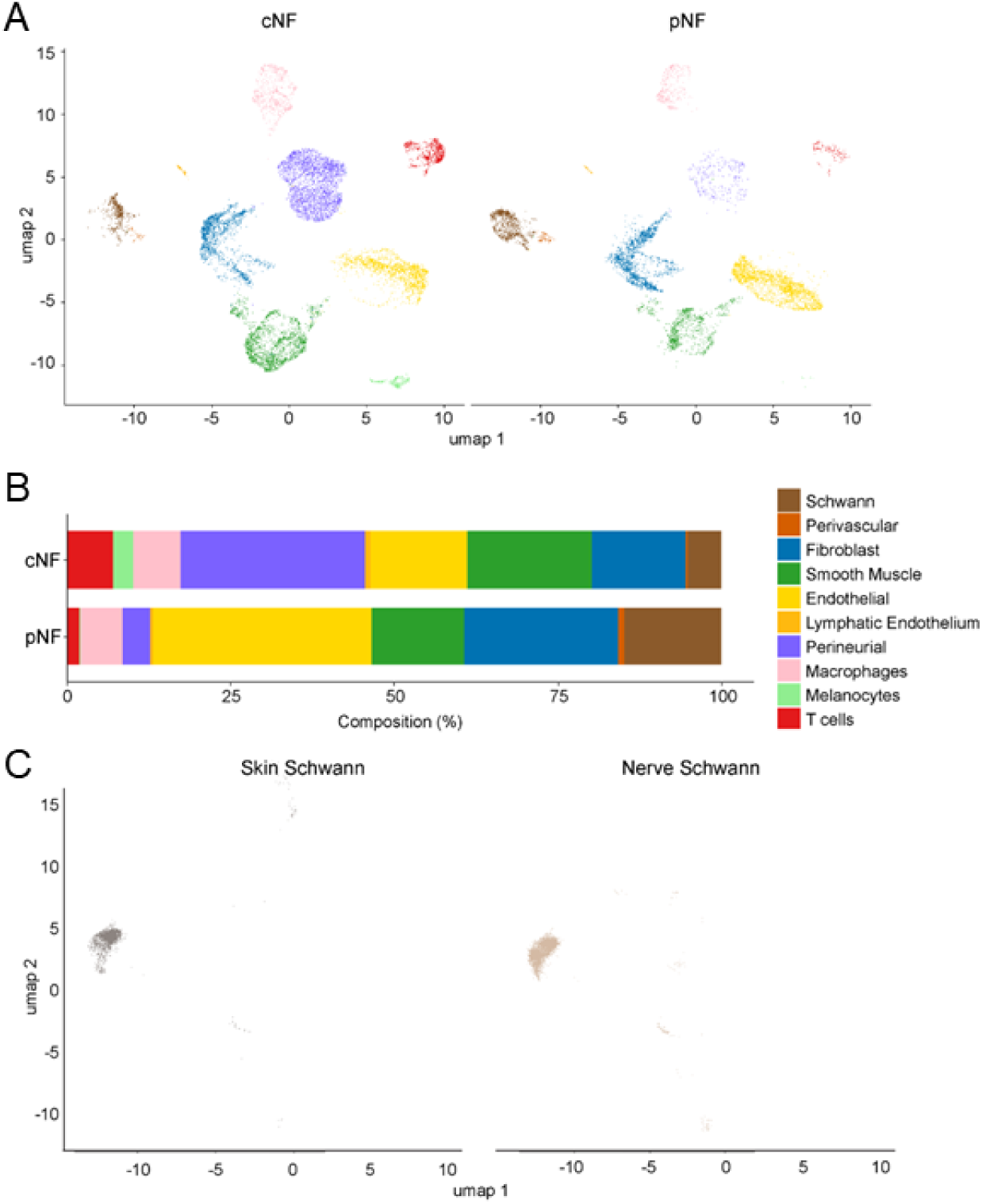
Cell type composition of cNF and pNF tumors compared with *NF1*-expressing reference Schwann cell populations. **(A)** UMAP visualization of single-cell transcriptomes from colored by annotated cell type. **(B)** Relative proportions of annotated cell types within cNF and pNF tumors. While both tumor types contain similar lineages, their cellular composition differs in the relative abundance of cell populations. **(C)** Projection of ***NF1*-expressing** skin- and nerve-derived Schwann cell references into the integrated UMAP space, showing alignment of each reference population with its tumor-associated Schwann cell states.

### 3.3 Subtype-Specific Transcriptional Programs in Schwann Cells Distinguish cNF from pNF tumors

Given the central role of Schwann cells in *NF1*-deficiency associated tumorigenesis, we next performed a focused comparison of their transcriptional states between cNF and pNF tumors. UMAP visualization highlighted distinct Schwann cell subpopulations associated with each tumor type (Figure 3A). Split UMAPs illustrate the distribution of Schwann cells by sample of origin, confirming alignment of skin-derived Schwann cells with cNF states and nerve-derived Schwann cells with pNF states (Supplemental Figure S2). To account for tissue-of-origin differences, differential expression was performed relative to *NF1*-expressing Schwann cells from matched skin or nerve sources. This analysis revealed both shared and unique transcriptional changes, with subsets of upregulated (Figure 3B) and downregulated (Figure 3C) genes distinguishing cNF- and pNF-derived Schwann cells (complete DEG lists are provided in Supplemental Table S2). A subset of genes was commonly altered across both tumor types, while others were enriched in only one context. Gene set enrichment analysis further revealed functional divergence (Figure S3), with pathways related to extracellular matrix organization and angiogenesis predominating in cNFs, while immune signaling and cell-cycle programs were more pronounced in pNFs. Expanded gene ontology enrichment results, grouped into refined functional categories, are provided in Supplemental Figure S3, with full enrichment outputs available in Supplemental Table S3. Selected genes of interest are highlighted in a dot plot (Figure 3D), emphasizing that although core Schwann lineage markers were maintained, *NF1* gene loss led to distinct biological programs shaped by tumor subtype.

**Figure 3.**
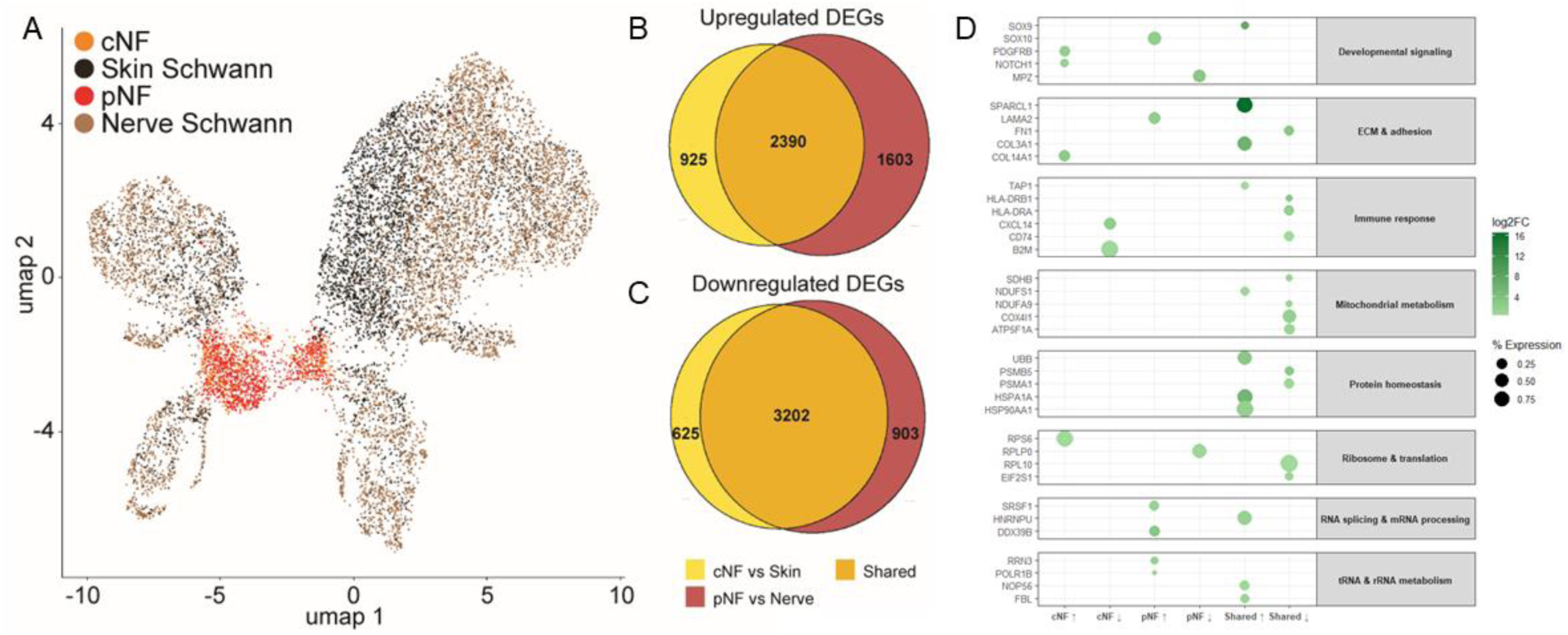
Differential expression analysis of Schwann cells highlights shared and subtype-specific transcriptional programs. **(A)** UMAP visualization of integrated Schwann cell populations from cutaneous neurofibromas (cNF), plexiform neurofibromas (pNF), and *NF1*-expressing skin- and nerve-derived references. **(B–C)** Venn diagrams showing the number of upregulated (B) and downregulated (C) differentially expressed genes (DEGs) in cNF versus skin Schwann cells and pNF versus nerve Schwann cells, with substantial overlap representing shared tumor-associated programs. **(D)** Dot plot summarizing representative DEGs grouped by functional categories, including developmental signaling, extracellular matrix (ECM) and adhesion, immune response, mitochondrial metabolism, protein homeostasis, ribosome and translation, mRNA processing, and rRNA metabolism. Dot size reflects the proportion of cells expressing each gene, while color intensity indicates average log2 fold-change.

In addition, to broad separation by tumor subtype and tissue of origin (Figure 3A), Schwann cells further resolved into multiple subclusters distinguished by unique transcriptional signatures (Supplemental Figure S2B). Marker analysis confirmed that these subclusters expressed distinct gene sets, including lineage-related factors (Supplemental Figure S2C). While a comprehensive analysis of Schwann heterogeneity was beyond the scope of the present study, we note that several subclusters showed preferential representation in either cNF or pNF tumors, suggesting that disease-specific programs identified by differential expression (Figure 3B–D) may be further stratified across discrete Schwann cell states.

### 3.4 Fibroblast analyses reveal tumor- and subtype-specific programs

To investigate how *NF1* gene loss alters fibroblast transcriptional states in neurofibromas, we compared fibroblast populations isolated from cutaneous (cNF) and plexiform (pNF) tumors to *NF1*-expressing dermal fibroblasts derived from normal skin (11, 21). UMAP visualization demonstrated clear separation of tumor-associated fibroblasts from the *NF1*-expressing dermal fibroblasts, with further divergence between cNF- and pNF-derived fibroblasts (Figure 4A, Supplemental Table S4). Differential expression analyses identified gene sets uniquely up- or down-regulated in each tumor subtype relative to dermal *NF1*-expressing fibroblasts (Figure 4B, Supplemental Table S5). The pNF tumoral fibroblasts comparison is relative to dermal fibroblasts (Supplemental Figure S4) and should be interpreted with this context in mind. Nevertheless, both comparisons revealed robust subtype-specific programs, and dot plot analysis highlighted canonical fibroblast markers and disease-associated genes distinguishing the three populations.

**Figure 4.**
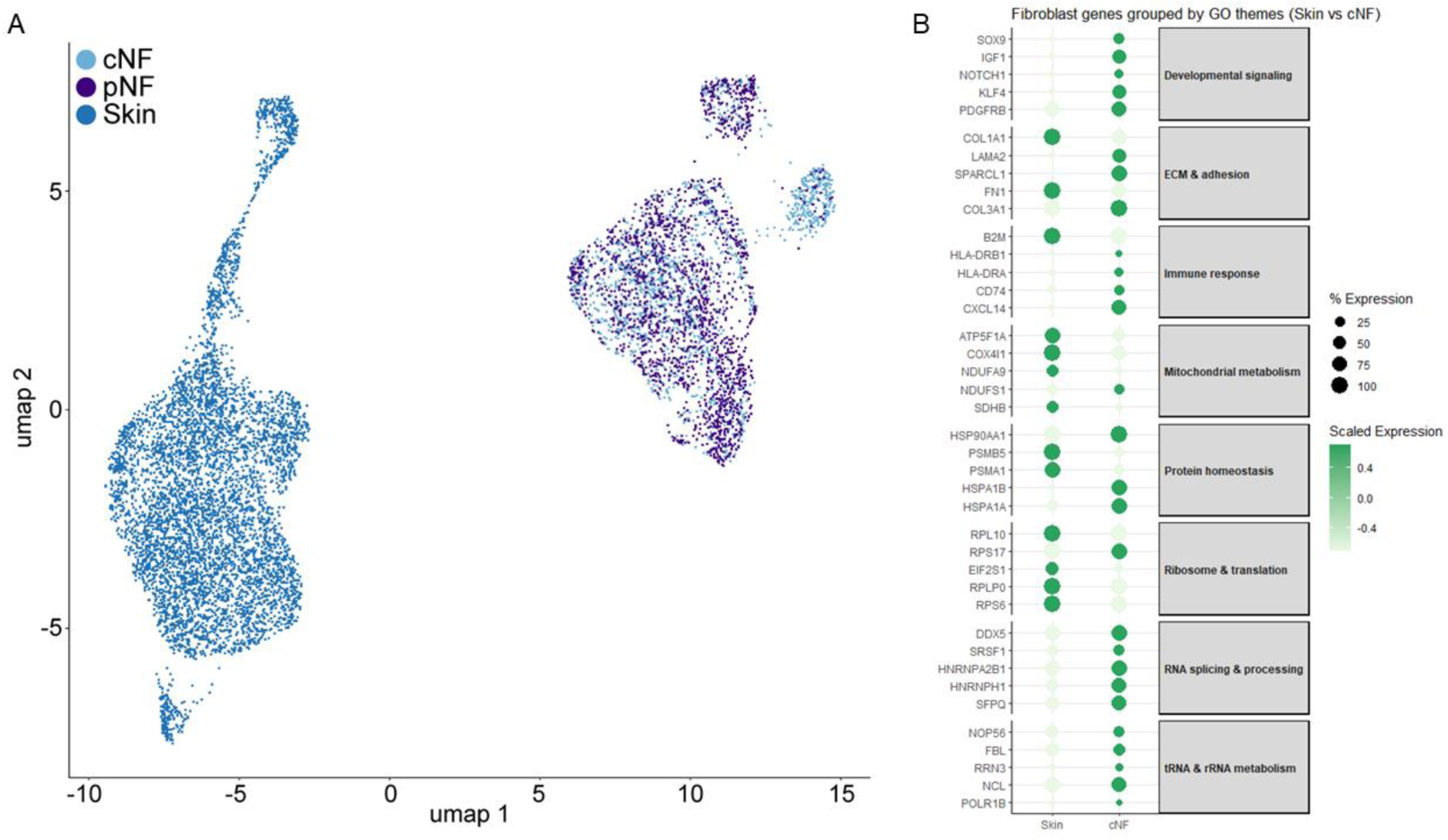
Tumor *NF1*-deficient fibroblast populations show transcriptional divergence from *NF1*-expressing dermal fibroblasts. **(A)** UMAP visualization of fibroblasts from cutaneous neurofibroma (cNF, light blue), plexiform neurofibroma (pNF, purple), and *NF1*-expressing dermal skin fibroblasts (dark blue), illustrating the distinct clustering of tumor-derived fibroblasts relative to their normal counterparts. **(B)** Dot plot showing expression of selected genes grouped by enriched gene ontology (GO) categories, comparing NF1-expressing skin fibroblasts to cNF fibroblasts. Dot size reflects the percentage of cells expressing each gene; color scale represents average scaled expression. Functional themes include developmental signaling, extracellular matrix (ECM) remodeling, immune response, mitochondrial metabolism, protein homeostasis, RNA processing, and translation-related programs. These features highlight tumor-associated reprogramming of fibroblasts in the context of *NF1* gene loss.

### 3.5 Distinct Extracellular Matrix and Growth Factor Signaling Networks Link Schwann Cells to the Neurofibroma Microenvironment

To define how Schwann cells integrate into the tumor microenvironment, we mapped ligand– receptor interactions in cNF and pNF tumors. Network plots revealed that Schwann cells act as central hubs, engaging most prominently with fibroblasts, perivascular cells, and endothelial lineages in both tumor types (Figure 5A–B, Supplemental Table S6).

**Figure 5.**
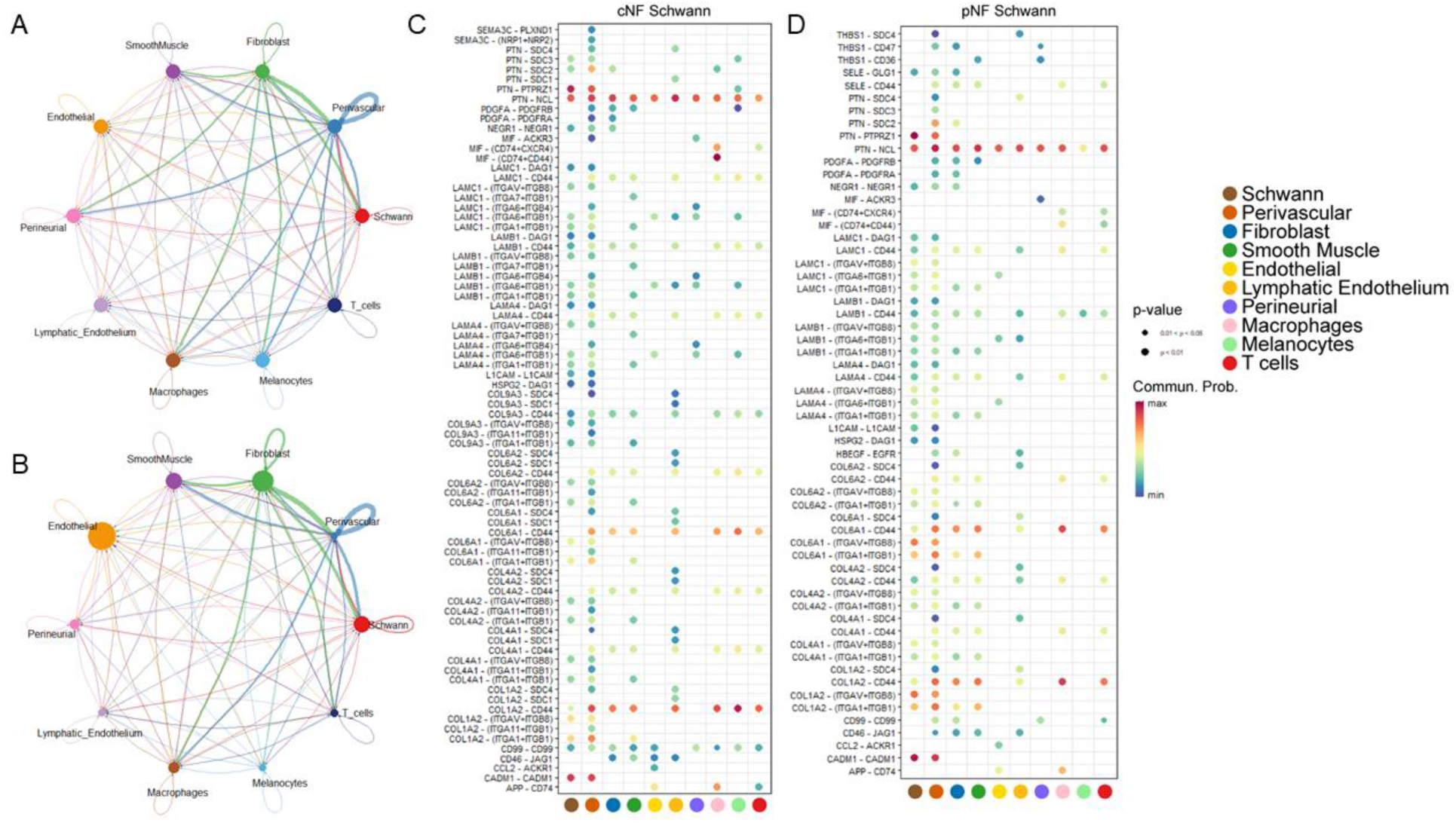
Cell–cell communication analysis of Schwann cells in cNF and pNF tumors. (A–B) Circle plots depicting the global communication networks of Schwann cells with other cell types in (A) cNF and (B) pNF tumors. Edge thickness is proportional to the communication probability, and colors represent the direction of signaling. (C–D) Dot plots showing ligand–receptor interactions involving Schwann cells in (C) cNF and (D) pNF. Each row corresponds to a ligand–receptor pair, and each column to a target cell type. Dot size reflects significance (p < 0.05 or p < 0.01), and dot color represents communication probability (red = higher, blue = lower). Cell type colors: Schwann (brown), Perivascular (orange), Fibroblast (blue), Smooth Muscle (green), Endothelial (yellow), Lymphatic Endothelium (light orange), Perineurial (purple), Macrophages (pink), Melanocytes (cyan), and T cells (red).

In cNF tumors, Schwann cells were enriched for adhesion- and growth factor–related signals, including *PTN–PTPRZ1*, *CADM1–CADM1*, and *NRXN1–NLGN1* interactions (Figure 5C). Multiple laminin–integrin pairs (e.g., *LAMA2–ITGA6*, *LAMB1–ITGA3*, and *LAMC1–ITGB1*) and collagen–integrin interactions (e.g., *COL6A2–ITGA1*, *COL14A1–ITGA2*) suggested robust Schwann–fibroblast and Schwann–perivascular crosstalk. *SPARCL1* also emerged as a recurrent Schwann-derived signal, linking to endothelial and stromal targets and reinforcing its role in modulating cell–matrix interactions. These pathways emphasize extracellular matrix remodeling and cell–cell adhesion as prominent features of the cNF microenvironment.

By contrast, pNF Schwann cells showed strong engagement of collagen- and laminin-mediated signaling, including *COL1A2–ITGA2*, *COL3A1–ITGB1*, *COL6A3–ITGA10*, and *LAMB2–ITGA3* (Figure 5D). Additional interactions such as *THBS1–CD47* and *HSPG2–SDC4* reflected enhanced stromal and endothelial connectivity. Compared to cNF tumors, *PTN*- and *CADM1*-driven signaling was reduced, whereas matrix-derived cues were more pronounced, suggesting distinct strategies of Schwann cell–directed remodeling in pNF tumors.

Across both tumor types, Schwann cells shared a set of conserved interactions, including *PDGFA– PDGFRA* and *NCAM1–FGFR1*, reinforcing their role as organizers of growth factor and adhesion signaling within neurofibromas. However, the relative prominence of *PTN*, *CADM1*, and *SPARCL1* in cNF tumor versus collagen- and laminin-centered pathways in pNF tumor underscores how tissue context influences the molecular wiring of Schwann cell communication.

Analysis of fibroblast-derived ligands revealed a core set of interactions that were shared between cNF and pNF tumors (Supplemental Figure S5, Supplemental Table S6). These included abundant extracellular matrix components such as laminins (*LAMC1, LAMB1, LAMB2, LAMA2, LAMA4*), collagens (*COL6A1/2/3, COL4A1/2, COL1A1/2*), and fibronectin (*FN1*), which engaged multiple stromal and immune cell types. Growth factors such as *VEGFA, PDGFD,* and *KITLG*, as well as signaling molecules including *PTN* and *JAG1*, were also broadly maintained across tumor types, suggesting a conserved fibroblast contribution to paracrine support of the tumor microenvironment. Although the overall interaction patterns were shared, the relative probability of signaling events was often stronger in pNF tumors, reflected by larger bubble sizes, particularly for collagen- and laminin-mediated interactions. These findings highlight fibroblasts as a central and consistent source of extracellular matrix and growth factor signals in both neurofibromas, with quantitative differences in signaling strength that may contribute to subtype-specific tumor architecture and cellular remodeling.

## 4. Discussion

This integrative single-cell analysis highlights how *NF1* gene loss reshapes Schwann cell identity in neurofibromas. By combining tumor-derived cNF- and pNF cells with skin- and nerve-derived *NF1*-expressing populations (11, 21), we were able to disentangle loss of *NF1*-driven transcriptional programs from tissue-of-origin effects. The emergence of distinct Schwann states that clearly separated tumoral from non-tumoral cells, and further distinguished cutaneous from plexiform neurofibromas, underscores the lineage-specific vulnerability of Schwann cells to *NF1* inactivation, a principle consistent with prior observations that Schwann cells are the cell of origin for *NF1*-associated peripheral nerve sheath tumors (27, 28). The ability to resolve these states within a unified atlas provides a framework for probing how *NF1*-associated alterations in Schwann cells and fibroblasts drive divergent tumor phenotypes and may help explain why clinically distinct neurofibroma subtypes arise in different tissue contexts (15, 29). Together, these findings establish a framework for probing how *NF1*-associated alterations in Schwann cells and fibroblasts drive divergent tumor phenotypes.

Building on the foundational characterization of neurofibromas (*NF1*-deficient) and *NF1*-expressing tissues by Wang et al. and Chu et al. respectively; our study integrates these datasets to directly investigate *NF1*-dependent transcriptional changes across Schwann cells and fibroblasts. By integrating these datasets, the present analysis extends their findings and reveals how tumor-associated Schwann cell states diverge from *NF1*-expressing reference populations. Whereas Wang et al. demonstrated the multicellular diversity of neurofibromas, including both Schwann and stromal populations, and Chu et al. characterized lineage-specific transcriptional programs in *NF1*-expressing skin and nerve, direct alignment of the two sources uncovered that Schwann cells from cNF and pNF tumors occupy distinct transcriptional niches relative to their skin- and nerve-derived counterparts. This integrative perspective highlights how tumor context interacts with developmental origin to shape Schwann cell states, an observation consistent with prior work suggesting that tissue environment contributes to *NF1*-deficient tumor heterogeneity (15, 29). Moreover, the finding that cNF- and pNF-derived Schwann cells retain transcriptional proximity to skin- and nerve-resident Schwann cells, respectively, provides evidence for origin-specific susceptibility that was not apparent in the individual datasets (28).

Subclustering further revealed heterogeneity within Schwann cell populations, identifying discrete regions enriched for extracellular matrix– and adhesion-related genes, including *SPARCL1*. The disease-associated expression of *SPARCL1* is notable given its established role in modulating cell– matrix interactions, angiogenesis, and tissue remodeling (39). Its localization to specific Schwann cell subsets suggests that *SPARCL1* may contribute to microenvironmental alterations unique to neurofibromas, particularly in shaping extracellular matrix composition and vascular responses. Importantly, *SPARCL1* also emerged in downstream ligand–receptor analyses, reinforcing its potential as a context-specific mediator of Schwann cell–stromal communication. Highlighting these recurrent disease-associated genes underscores how *NF1* gene loss drives not only broad lineage programs but also focal adaptations within Schwann cell subpopulations that may carry functional significance.

Projecting *NF1*-expressing Schwann cells and fibroblasts onto the neurofibroma (*NF1*-deficient) atlas further clarified how tissue origin and *NF1* status interact to shape transcriptional programs. Skin-derived Schwann cells aligned closely with Schwann cell clusters from neurofibromas, suggesting that cNF-associated states may reflect adaptations of skin-resident lineages that normally contribute to sensory function and cutaneous homeostasis (30). In contrast, nerve-derived Schwann cells mapped to a distinct subset, consistent with the idea that pNF-associated programs arise from context-specific vulnerabilities of nerve-associated Schwann cells, which are known to activate specialized repair and proliferation programs following injury (31–33). Similarly, dermal fibroblasts projecting onto the tumor fibroblast populations, confirms lineage fidelity while also highlighting transcriptional divergence that emerges in the *NF1*-deficient setting, aligning with prior observations that fibroblasts actively remodel extracellular matrix and influence Schwann cell behavior in the tumor microenvironment (34–36). Together, these projections demonstrate that neurofibroma cell populations encompass canonical cell identities while simultaneously acquiring *NF1*-gene-loss–dependent adaptations, enabling direct comparison between *NF1*-expressing and *NF1*-deficient tumor contexts.

Differential expression analysis revealed specific markers that distinguish *NF1*-deficient Schwann cells from *NF1*-expressing reference populations. In particular, *NRXN1*, *GALR1*, and *COL28A1* were consistently upregulated in both cNF- and pNF-derived Schwann populations relative to *NF1*-expressing skin- and nerve-derived controls, highlighting a set of shared disease-associated transcriptional features. These markers suggest alterations in cell–cell adhesion (*NRXN1*) (37, 38), G-protein–coupled signaling (*GALR1*) (39, 40), and extracellular matrix organization (*COL28A1*) (41, 42), pointing to convergent pathways that may support tumor growth and maintenance. *NF1*-expressing Schwann cells were enriched for transcripts including *APOD, DCN, MDK, FOSB, SPARCL1, ERBB3*, and *COL5A3*, which encompass regulators of lipid metabolism (*APOD*) (43), extracellular matrix organization (*DCN, COL5A3*) (44–46), growth factor signaling (*MDK, ERBB3*) (47–49), and activity-dependent transcription (*FOSB*) (50). These lineage-associated programs underscore functional differences that are attenuated or lost following *NF1* inactivation, reflecting transcriptional adaptations driven by *NF1* gene loss rather than tissue-of-origin effects.

Building on these gene-level differences, focused analysis of Schwann cell populations demonstrated that *NF1* gene loss induces both shared and subtype-specific transcriptional programs. While lineage-defining markers such as *SOX10* and *PLP1* remained intact, differential expression analysis revealed broad remodeling of pathways governing cellular function. Genes involved in extracellular matrix organization and angiogenic signaling were preferentially upregulated in cNF-derived Schwann cells, consistent with reports that cutaneous tumors exhibit increased stromal expansion and vascular remodeling (51). In contrast, pNF-derived Schwann cells showed enrichment of immune response and cell-cycle–associated pathways, in line with prior evidence that plexiform lesions are characterized by heightened macrophage infiltration, inflammatory cytokine signaling, and proliferative potential (32, 44). The presence of a substantial set of genes commonly altered across both cNF and pNF tumors indicates conserved effects of *NF1* inactivation, whereas the subtype-specific signatures highlight context-dependent adaptations. Together, these findings suggest that while *NF1* gene loss establishes a core Schwann cell disease program, tumor subtype and tissue of origin direct additional transcriptional trajectories that may underlie clinical differences between cutaneous and plexiform neurofibromas.

Projecting *NF1*-expressing reference cells onto the neurofibroma (*NF1*-deficient) atlas further underscored how tissue origin interacts with *NF1* status to shape cellular programs. Skin-derived Schwann cells localized near cNF states, whereas nerve-derived Schwann cells aligned with pNF states, confirming that developmental origin imprints remain evident even after *NF1* gene loss. Dermal fibroblasts similarly mapped into tumor fibroblast clusters, validating their lineage fidelity while highlighting transcriptional divergence that arises in the *NF1*-deficient context. At the tissue level, comparisons of cellular composition revealed both shared diversity and subtype-specific biases. cNF tumors were enriched for perineurial and T cells, consistent with a previous study showing cNF-intratumoral T cells (52), whereas pNFs were dominated by Schwann cell, fibroblast and perivascular lineages, reflecting their nerve-associated niche and characteristic hypervascularity. Together, these projections and compositional shifts provide a mechanistic framework in which *NF1* inactivation overlays lineage-specific programs, producing distinct tumor ecologies that align with the clinical and histological features of cutaneous and plexiform neurofibromas.

Among the most prominent Schwann cell markers identified in this analysis was *PTN* (pleiotrophin), which was strongly upregulated in both cNF- and pNF-derived populations compared to *NF1*-expressing Schwann cells. *PTN* was expressed in the majority of tumor Schwann cells (∼61% in cNFs; ∼66% in pNFs) versus a small minority of their *NF1*-expressing skin- and nerve-derived counterparts, with nearly seven-fold higher abundance in cNF tumors and robust increases in pNF tumors (log₂FC = 6.97 and 6.47, respectively; adj. p < 0.01). Beyond serving as a consistent disease-associated marker, CellChat analysis identified *PTN* as a secreted ligand engaging *NCL* (nucleolin), a receptor that regulates ribosome biogenesis, chromatin structure, and rRNA processing. *PTN–NCL* interactions are known to activate *MAPK* and *PI3K/AKT* signaling and have been implicated in angiogenesis, extracellular matrix remodeling, and tumor–stroma communication, suggesting a potential mechanism by which *PTN* contributes to neurofibroma pathogenesis. Similarly, *CADM1* (cell adhesion molecule 1) was consistently upregulated across both tumor types (log₂FC = 2.26 in cNFs; 2.86 in pNFs; adj. p < 0.01), expressed in ∼65–66% of the tumoral Schwann cells. As a mediator of axon–glia interactions, synaptic organization, and cell survival, *CADM1* upregulation likely reflects remodeling of adhesive and signaling interactions in *NF1*-deficient Schwann cells, with implications for both intrinsic cell behavior and cross-talk with stromal and neural elements. By contrast, *COL1A2*, despite its role in extracellular matrix biology, was not significantly altered in Schwann cells from either cNF or pNF tumors relative to *NF1*-expressing controls, indicating that its expression is not directly influenced by *NF1* gene loss. Together, these findings highlight *PTN* and *CADM1* as robust and functionally relevant markers of *NF1*-deficient Schwann cells, while also illustrating that not all extracellular matrix–associated genes are subject to *NF1*-dependent regulation. In contrast to *PTN-COL1A2* was not significantly altered in Schwann cells from either cNF or pNF tumors compared to *NF1*-expressing controls. This indicates that *COL1A2* expression is not directly influenced by *NF1* gene loss within Schwann cells.

Beyond transcriptional markers, comparisons of cellular composition between cNF and pNF tumors revealed both shared lineage diversity and subtype-specific biases. The enrichment of perineurial, endothelial, and fibroblast populations in cNF tumors suggests that microenvironmental remodeling and stromal expansion are central features of cutaneous tumors, consistent with prior histological descriptions of dense collagen deposition and vascular proliferation in cNF tumors (15, 29). By contrast, the predominance of Schwann and perivascular lineages in pNF tumors underscores the nerve-associated niche and its supportive vasculature as defining aspects of plexiform disease, in agreement with reports of hypervascularity and Schwann cell expansion in these lesions (53, 54). Importantly, mapping of *NF1*-expressing Schwann cells confirmed that these differences align with developmental origin, as *NF1*-expressing skin-derived Schwann cells localized near cNF states while nerve-derived Schwann cells clustered with pNF states. These observations suggest that both tissue of origin and *NF1* inactivation cooperate to shape tumor subtype–specific transcriptional programs, offering a mechanistic explanation for the distinct biological and clinical behavior of cNF and pNF tumors.

In addition to Schwann cell alterations, our analysis highlighted transcriptional adaptations in fibroblasts that contribute to the neurofibroma microenvironment. While dermal fibroblasts projected into tumor fibroblast clusters, confirming lineage fidelity, *NF1*-deficient fibroblasts exhibited distinct signatures characterized by upregulation of extracellular matrix–associated transcripts. This is consistent with prior reports that fibroblasts in neurofibromas actively remodel matrix components and influence Schwann cell behavior through paracrine signaling (55, 56). The expansion of fibroblast populations observed in cNF tumors further supports their role in driving stromal deposition and vascular remodeling, hallmarks of cutaneous lesions. These findings underscore that *NF1* gene loss reprograms not only Schwann cells but also fibroblasts, and that reciprocal interactions between these two lineages may be central to shaping the tumor niche.

In summary, this integrative single-cell analysis reveals how *NF1* gene loss reshapes both Schwann cells and fibroblasts, establishing a shared core disease program while allowing tissue-of-origin and tumor context to direct additional subtype-specific trajectories. The identification of recurrent markers such as *PTN, CADM1, NRXN1, GALR1, COL28A1* and *SPARCL1*, along with pathway-level distinctions between cNF- and pNF-derived Schwann cells, underscores the multifaceted ways in which *NF1* inactivation drives tumor biology. Importantly, by aligning *NF1*-deficient tumor-derived populations with *NF1-*expressing counterparts, our study disentangles *NF1*-dependent transcriptional adaptations from lineage-intrinsic programs, providing a mechanistic framework to explain the distinct cellular composition, stromal remodeling, and clinical behavior of cutaneous versus plexiform neurofibromas.

## 5. Supplemental Figures and Tables

**Supplemental Figure S1.**
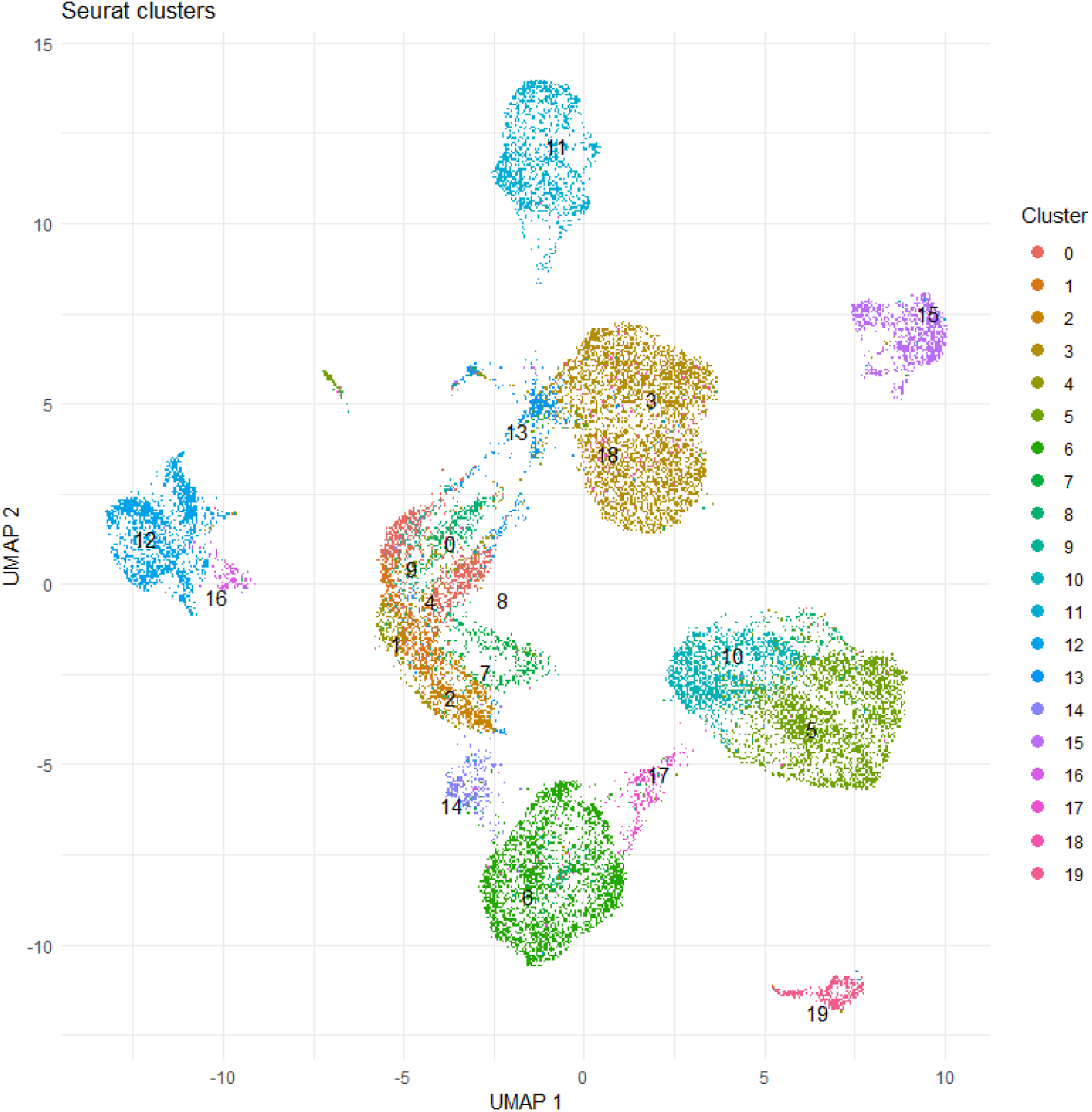
Unsupervised clustering of integrated single-cell transcriptomes. UMAP embedding of all cells following integration and normalization, colored by Seurat-derived clusters (0–19). Each unbiased cluster represents a transcriptionally distinct cell population prior to annotation.

**Supplemental Table S1. Marker genes for Seurat clusters.** List of significantly enriched genes for each cluster identified in the integrated single-cell dataset. Shown are cluster assignment, gene symbol, average log fold-change, percentage of *NF1*-expressing cells in the cluster versus all other cells, and adjusted *p*-values. These markers were used to guide cell type annotation.

**Supplemental Table S2. Differentially expressed genes in Schwann cells.** Upregulated (Figure 3B) and downregulated (Figure 3C) genes distinguishing cNF- and pNF-derived Schwann cells. Complete gene lists are provided here.

**Supplemental Table S3. Gene set enrichment analysis of Schwann cell transcriptional programs.** Gene set enrichment analysis revealed functional divergence between cNF and pNF Schwann cells. Expanded enrichment results, grouped into refined functional categories, are summarized in Supplemental Figure S3, with the complete outputs provided here.

**Supplemental Figure S2.**
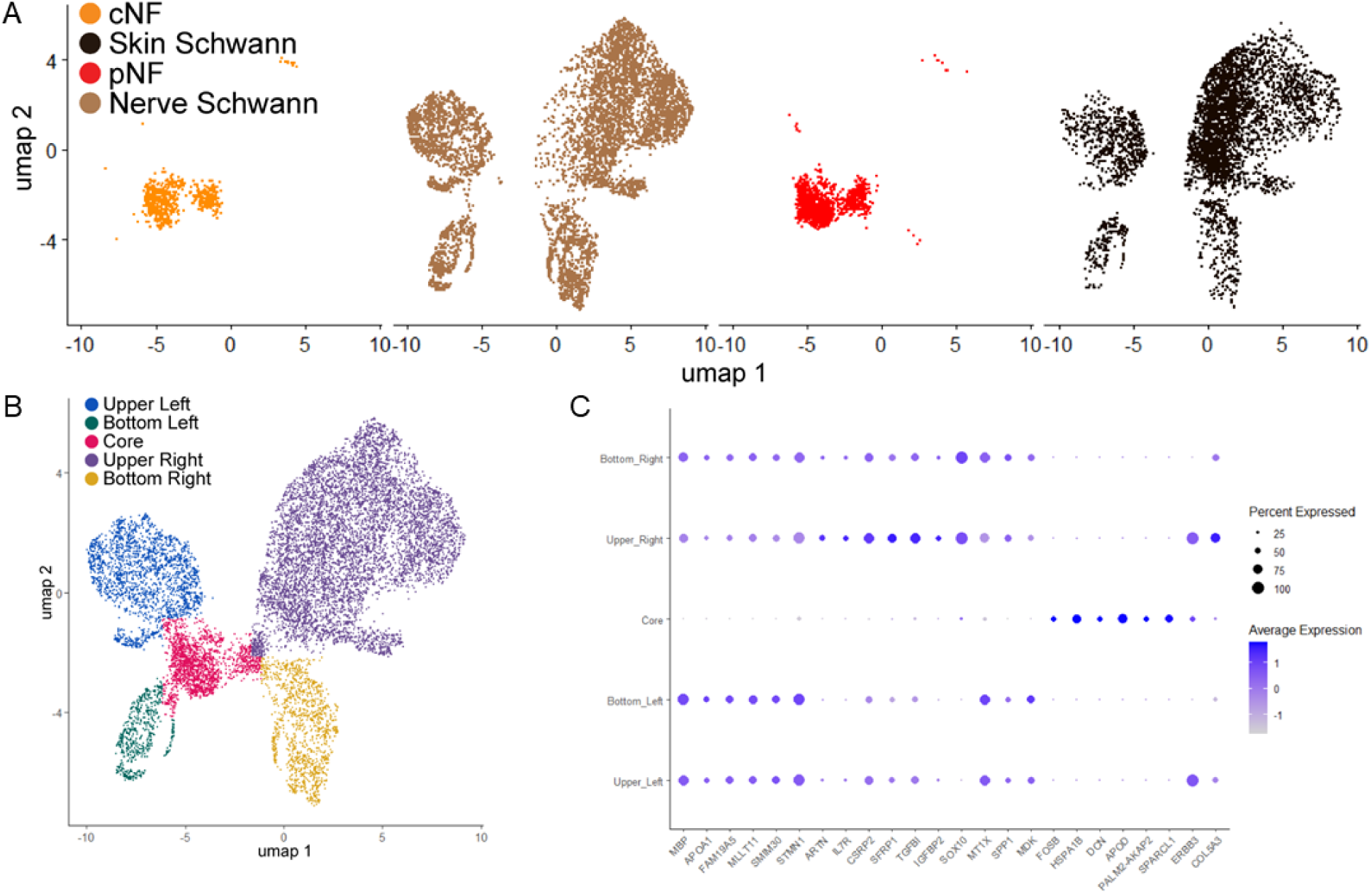
Integration and regional characterization of Schwann cells from neurofibromas and *NF1*-expressing tissues. (A) UMAP visualization of integrated single-cell RNA-seq datasets showing Schwann cells derived from cNF, (orange), pNF (red), nerve (brown), and skin (black). (B) UMAP of the integrated dataset with five spatially distinct Schwann cell regions: Upper Left (blue), Bottom Left (green), Core (pink), Upper Right (purple), and Bottom Right (yellow). (C) Dot plot showing representative marker genes for each Schwann cell region.

**Supplemental Figure S3.**
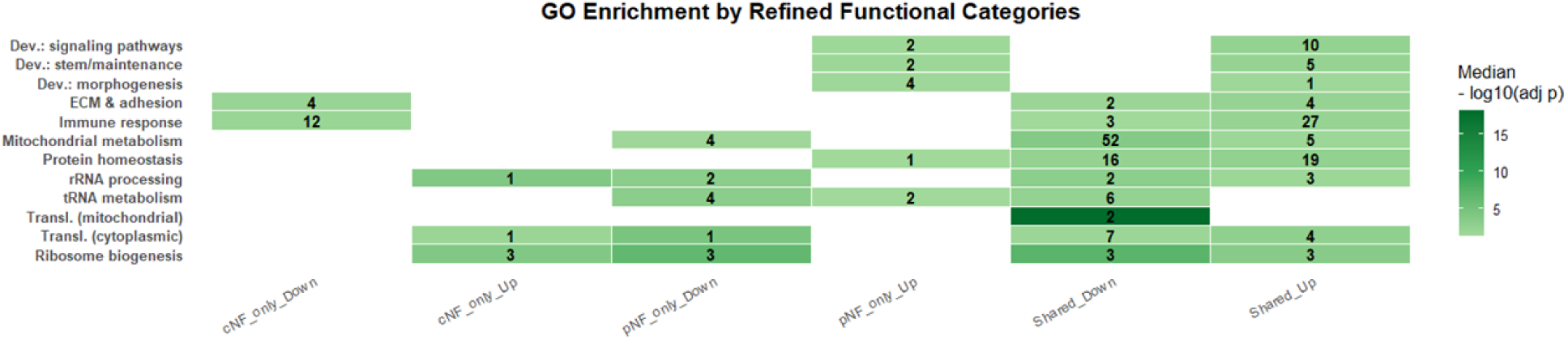
Unique enriched pathways in cNF- or pNF-derived Schwann cells. Heatmap showing enriched pathways for up- and down-regulated gene sets unique to cNF- or pNF-derived Schwann cells, as well as those shared across tumor types. Numbers within boxes denote the count of significant terms per category; color intensity corresponds to the median −log10(adjusted *p*-value).

**Supplemental Figure S4.**
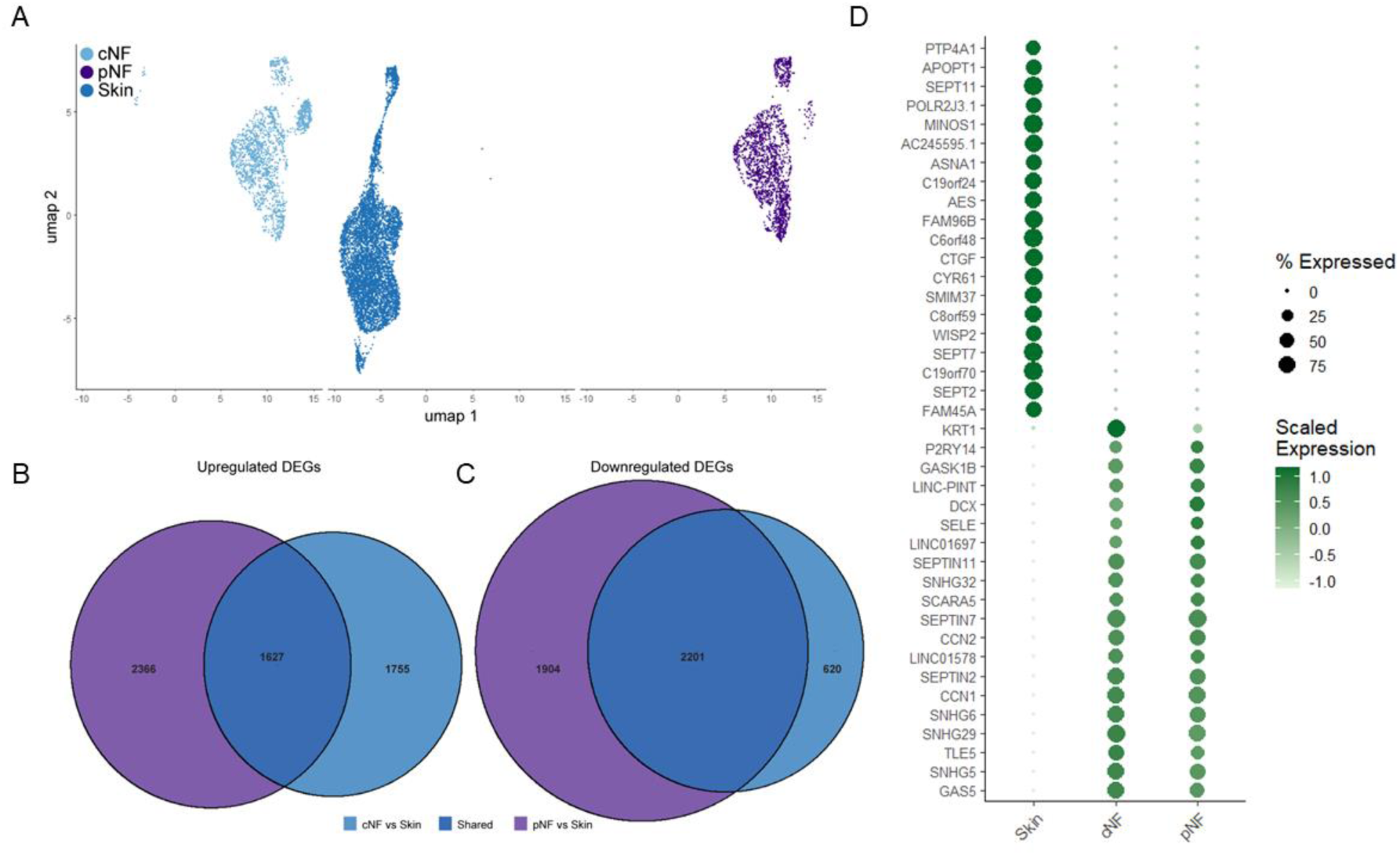
Tumor-specific and shared transcriptional changes in fibroblasts from cNF and pNF tumors. (A) UMAP projection of fibroblast populations derived from cNF, pNF, and *NF1*-expressing skin. (B) Venn diagram showing the number of significantly upregulated genes in cNF vs. skin (purple) and pNF vs. skin (blue) fibroblasts. (C) Venn diagram of downregulated genes in cNF vs. skin and pNF vs. skin fibroblasts. (D) Dot plot displaying the scaled expression and percent of cells expressing selected shared and condition-specific marker genes across skin-, cNF-, and pNF-derived fibroblasts. Dot size corresponds to the percentage of cells expressing each gene, and color intensity reflects scaled average expression.

**Supplemental Table S4. Differentially expressed genes in fibroblasts.** Differential expression analyses identified gene sets uniquely up- or down-regulated in each tumor subtype relative to dermal *NF1*-expressing fibroblasts (Figure 4B). Complete gene lists are provided here.

**Supplemental Table S5. Gene set enrichment analysis of fibroblast transcriptional programs.** Gene set enrichment analysis revealed functional divergence between cNF and pNF fibroblasts. Complete outputs provided here.

**Supplemental Figure S5.**
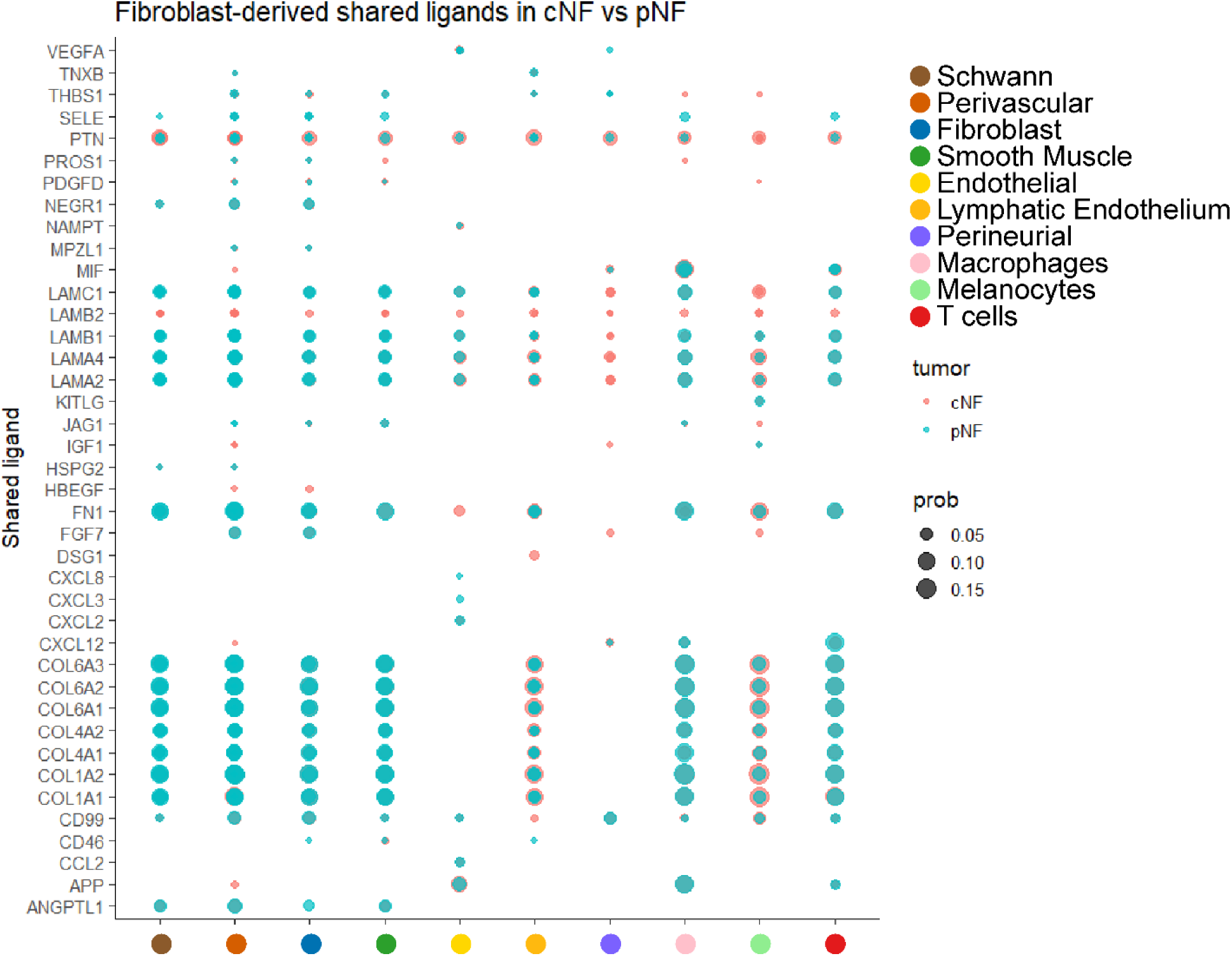
Fibroblast-derived shared ligands in cNF vs pNF tumors. Bubble plot showing fibroblast-derived ligands detected in both cNF (red) and pNF (teal) tumors. Each row represents a shared ligand, and each column corresponds to the interacting target cell type. Bubble size reflects the communication probability, with larger circles indicating higher interaction strength.

**Supplemental Table S6. Cell ligand–receptor interactions in neurofibromas.** CellChat analysis of cNF and pNF tumors. Complete receptor–ligand interaction lists are provided here.

